## Supplementary informations for "Molecular and evolutionary basis of O-antigenic polysaccharide driven phage sensitivity in environmental pseudomonads"

**Running title:** O-antigen-mediated phage and *Pseudomonas* interaction

Jordan Vacheron[1#], Clara M. Heiman[1], Julian R. Garneau[1], Peter Kupferschmied[1†], Ronnie de Jonge[2], Daniel Garrido-Sanz[1], Christoph Keel[1#]

[1]: Department of Fundamental Microbiology, University of Lausanne, Lausanne, Switzerland.
[2]: Plant-Microbe Interactions, Department of Biology, Science4Life, Utrecht University, Padualaan 8, Utrecht, 3584 CH, The Netherlands.

[†]: Present address: Peter Kupferschmied, Federal Office for Agriculture, Swiss Federal Plant Protection Service, Bern, Switzerland

### Content:

#### Supplementary figures

**Fig. S1.** Distribution of the transposon insertions across the genome of *Pseudomonas protegens* CHA0.
**Fig. S2.** Biological reproducibility of the Tn-seq results.
**Fig. S3.** Clusters of orthologous groups (COGs) classifications of the genes potentially involved in phage resistance according to the multiplicity of infection (MOI) conditions.
**Fig. S4.** Comparison of the Tn-seq data for the different phage concentrations tested.
**Fig. S5.** Impact of multiple and individual gene deletions within the OBC3 gene cluster of *Pseudomonas protegens* CHA0 on its phage susceptibility.
**Fig. S6.** Prediction and protein structure comparison of the putative pectate lyase encoded in the genome of the phage ΦGP100.
**Fig. S7.** Different sensitivity patterns displayed by *Pseudomonas protegens* subgroup strains that harbor the OBC3 gene cluster.
**Fig. S8.** The OBC3 gene cluster is located within a potential genomic island.
**Fig. S9.** Horizontal gene transfer signature detected within the different orthologous genomic regions of OBC3 in bacterial genomes.
**Fig. S10.** Maximum likelihood trees of the amino acid sequences of the corresponding genes conserved within the OBC3 gene cluster detected in bacterial genomes.

#### Supplementary tables

**Table S1**. Plasmids used in this study.
**Table S2.** Tn-seq characteristics.
**Table S3:** Phage genomes used in this study.
**Table S4.** Update of the phage GP100 annotation.
**Table S5.** Bacterial strains used in this study.
**Table S6.** Occurrence of inverted repeats sequences inside the OBC3 gene cluster of *Pseudomonas protegens* CHA0.
**Table S7**. *Pseudomonas protegens* CHA0 derivatives and *Escherichia coli* strain used in this study.
**Table S8**. Primers used in this study.

#### References

#### Supplementary figures

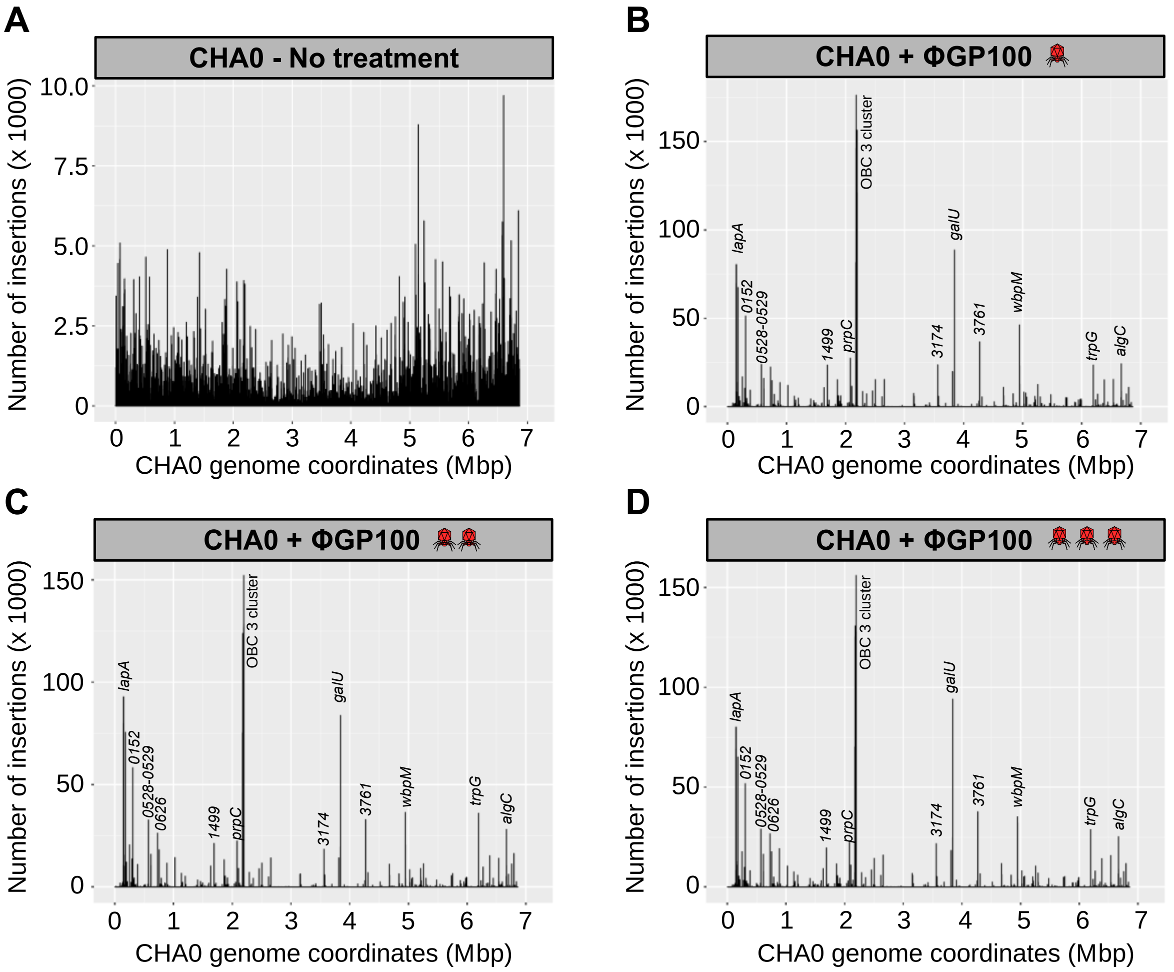


**Fig. S1. Distribution of the transposon insertions across the genome of *Pseudomonas protegens* CHA0**. Transposon insertions in the non-treated condition (**A**) and in presence of the phage ΦGP100 with a multiplicity of infection (MOI) of 1 (**B**), 10 (**C**) and 100 (**D**), indicated with cartoons of one, two or three red phages, respectively.

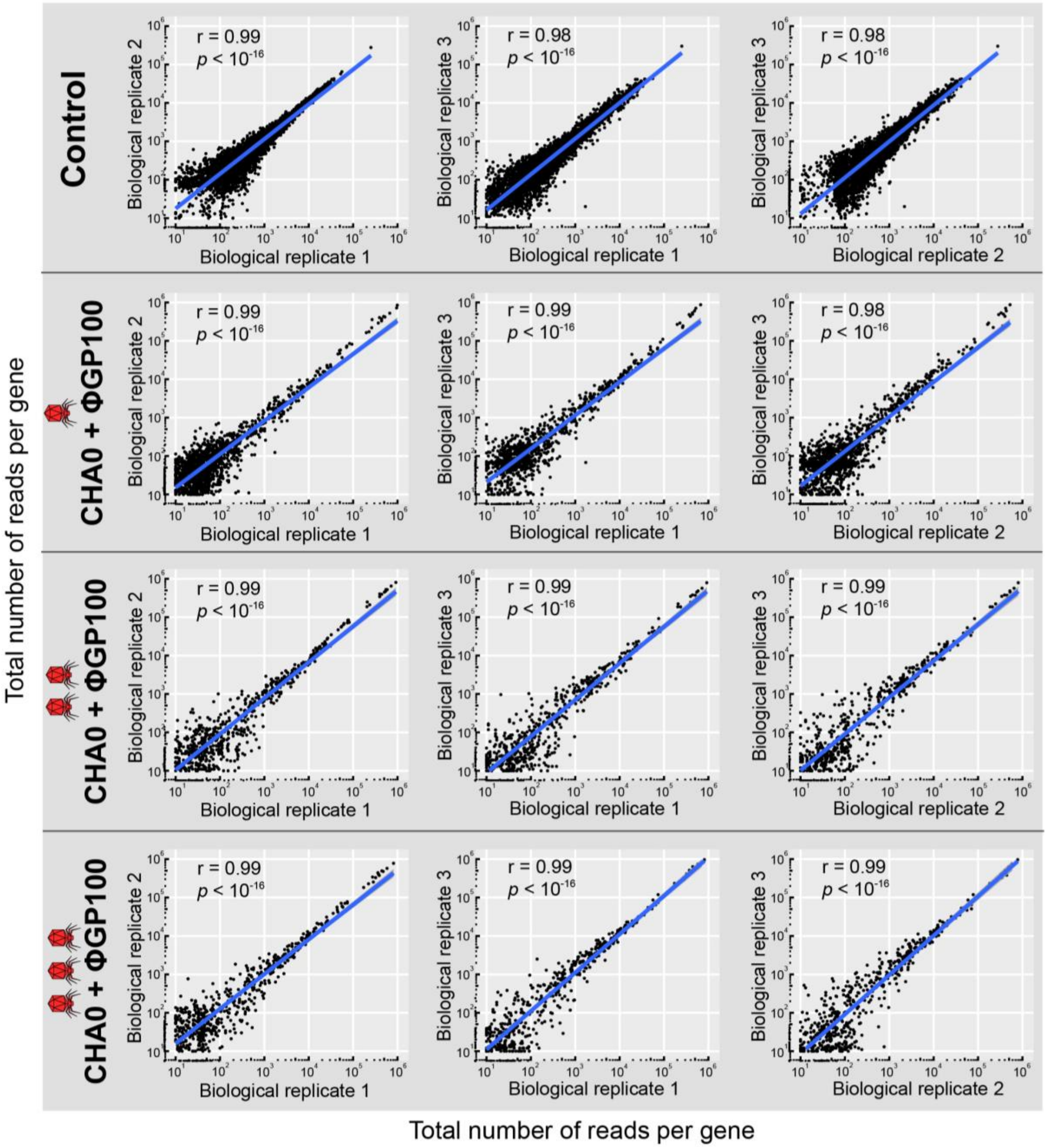


**Fig. S2. Biological reproducibility of the Tn-seq results.** Axes represent the total number of reads per gene for the respective biological replicates. Three independent biological replicates were compared for each condition. The different multiplicity of infection (MOI) conditions are represented with the red phage cartoons. One phage, MOI of 1; two phages, MOI of 10; three phages, MOI of 100. Control, no phage treatment. Correlation coefficients were calculated using Pearson correlation test.

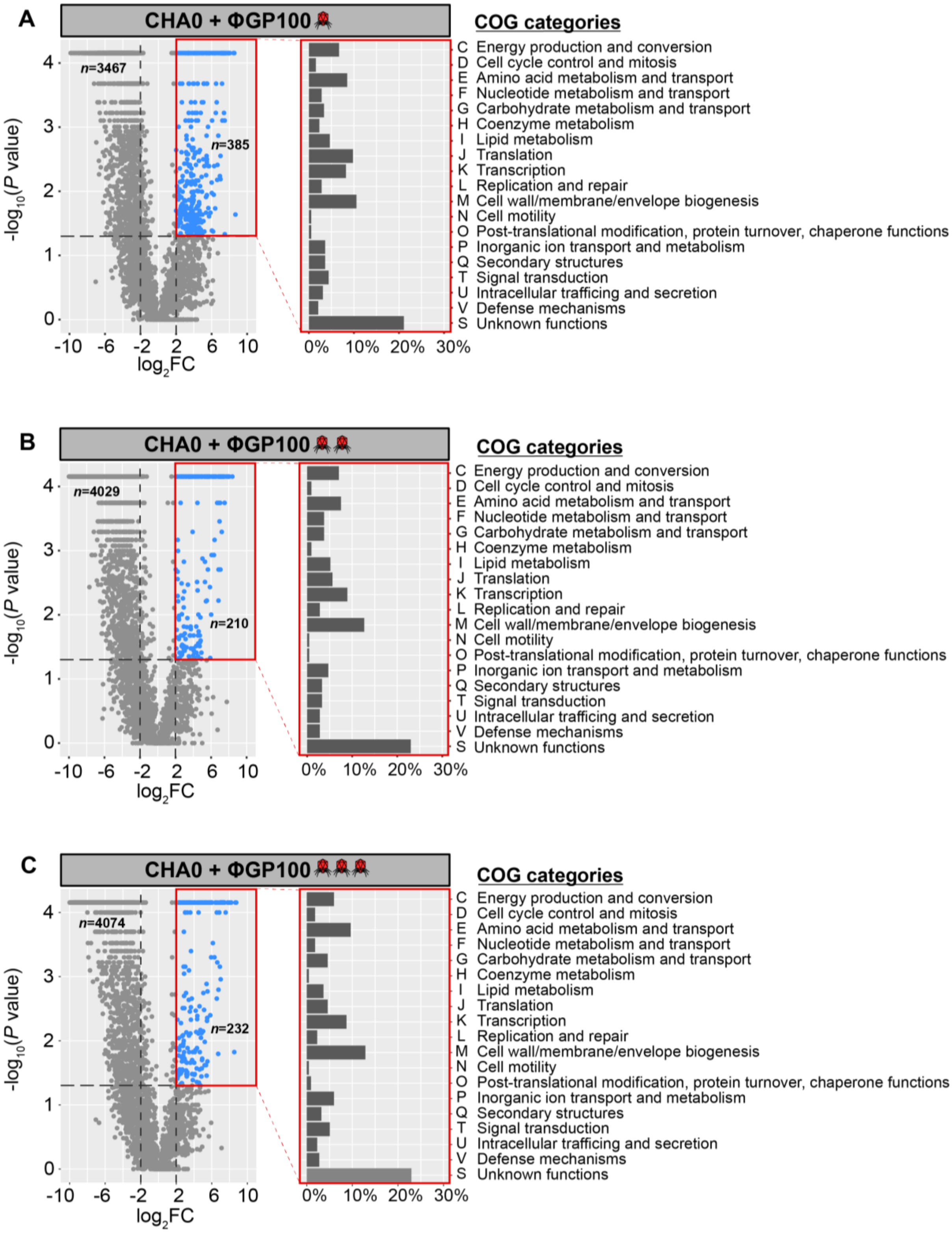


**Fig. S3. Clusters of orthologous groups (COGs) classifications of the genes potentially involved in phage resistance according to the multiplicity of infection (MOI) conditions.** Red phage cartoons: one phage, MOI of 1 (**A**); two phages, MOI of 10 (**B**); three phages, MOI of 100 (**C**). The blue dots correspond to genes presenting a log2FC > 2 and *p* < 0.05.

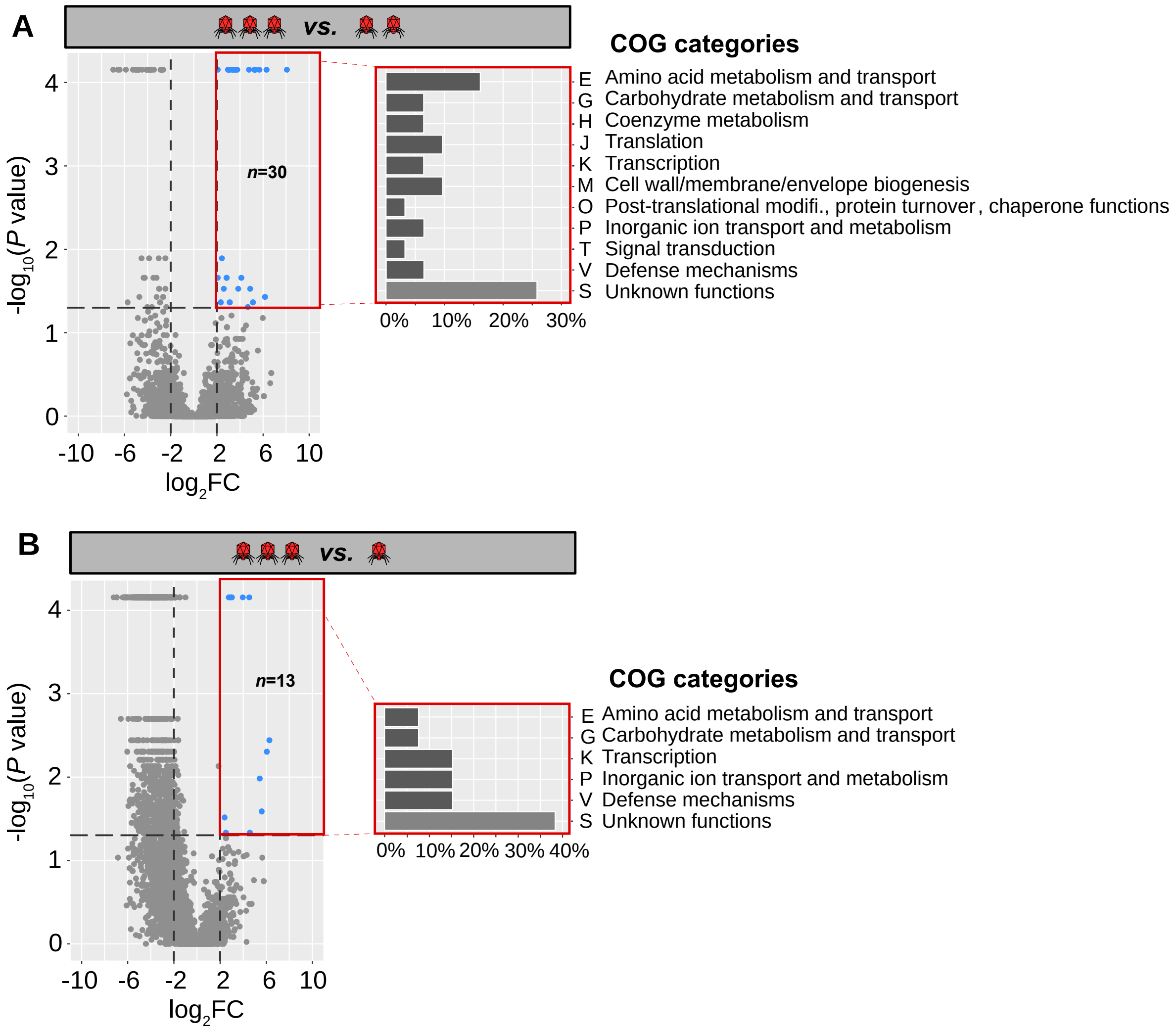


**Fig. S4. Comparison of the Tn-seq data for the different phage concentrations tested.** (**A**) Multiplicity of infection (MOI) of 100 vs. MOI of 10 and (**B**) MOI of 100 vs. MOI of 1. Red phage cartoons: one phage, MOI of 1; two phages, MOI of 10; three phages, MOI of 100. The genes potentially required for phage resistance are presented with blue dots (log2FC > 2 and $p < 0.05$) and classified according to the clusters of orthologous groups (COGs) they belong to.

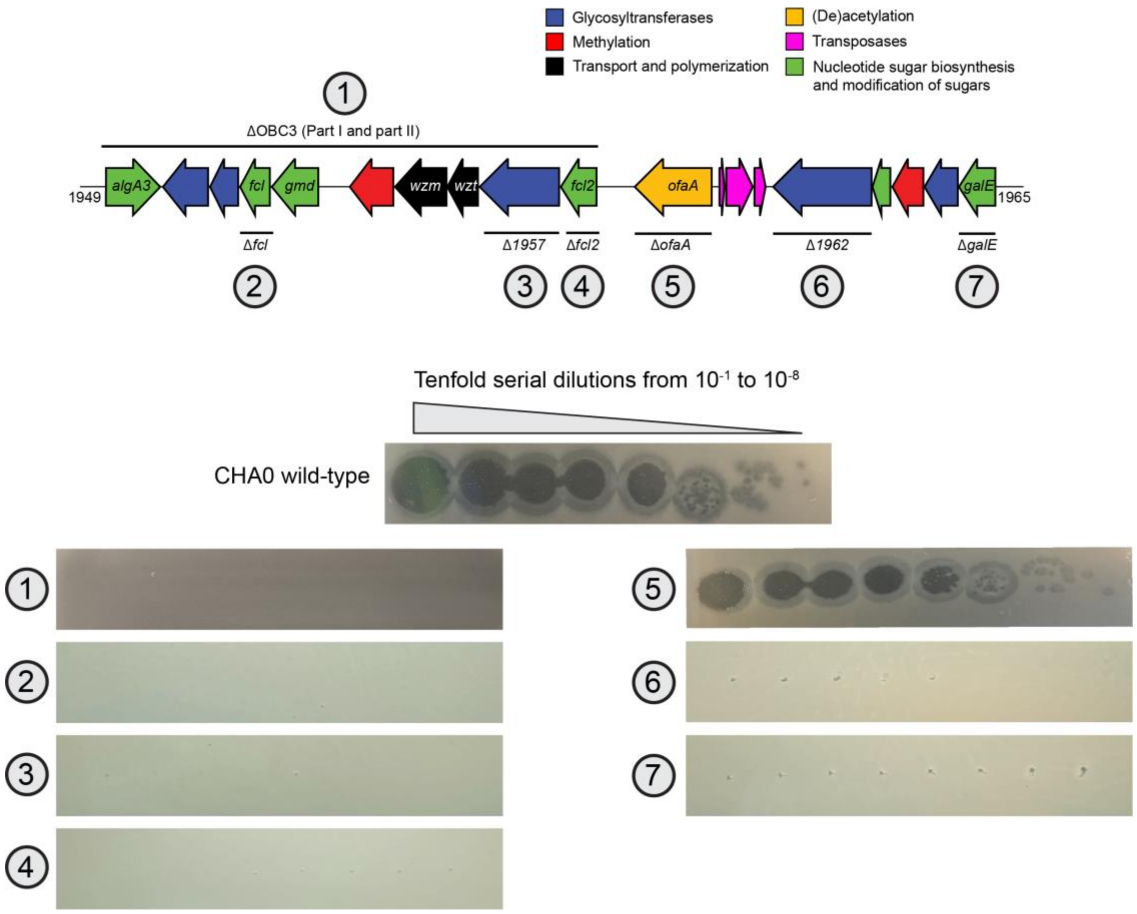


**Fig. S5. Impact of multiple and individual gene deletions within the OBC3 gene cluster of *Pseudomonas protegens* CHA0 on its phage susceptibility.** Spot assays of the phage ΦGP100 onto a bacterial lawn of CHA0 wild type or mutants. The deletion of the *ofaA* gene encoding an O-antigen acetylase has no influence on the sensitivity of CHA0 towards ΦGP100.

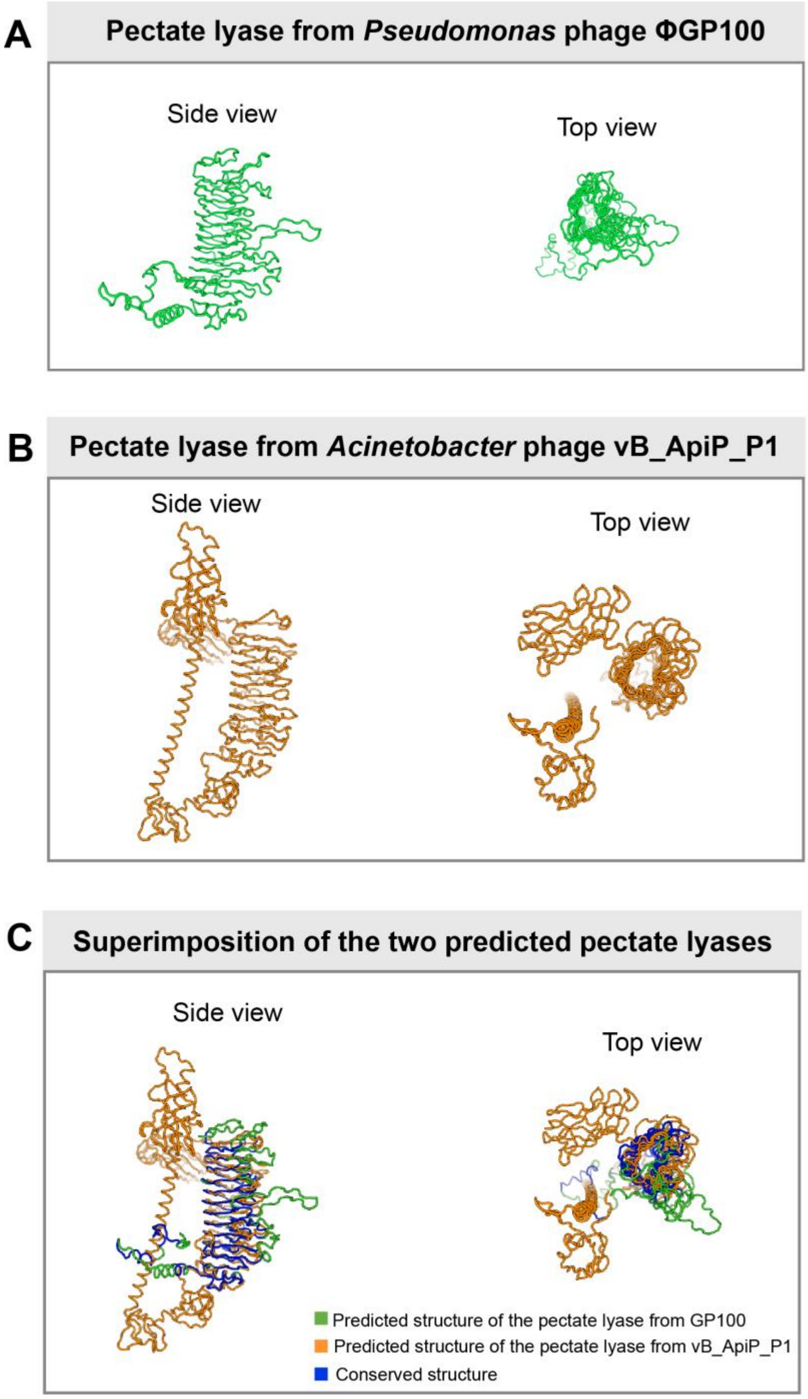


**Fig. S6. Prediction and protein structure comparison of the putative pectate lyase encoded in the genome of the phage ΦGP100. (A)** AlphaFold2 prediction of the protein structure of the pectate lyase of the phage ΦGP100.**(B)** AlphaFold2 prediction of the protein structure of the pectate lyase of the phage vB_ApiP_P1 previously demonstrated to display a depolymerase activity towards *Acinetobacter* cells (1). **(C)** Superimposition of the two pectate lyases shows a conserved structure.

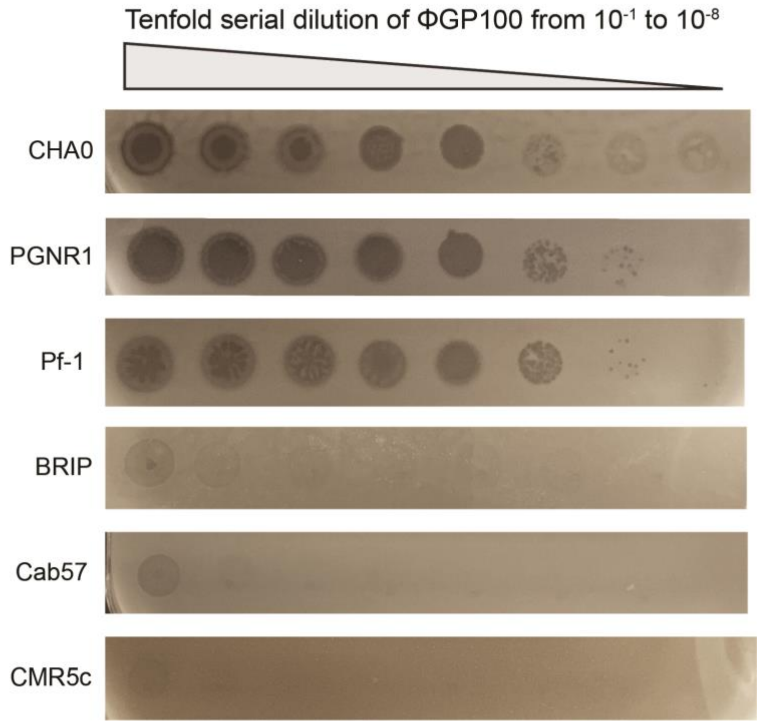


**Fig. S7. Different sensitivity patterns displayed by *Pseudomonas protegens* subgroup strains that harbor the OBC3 gene cluster.** Spot assays of ΦGP100 onto bacterial lawns of the different *Pseudomonas* strains. Strains CHA0, PGNR1 and Pf1 exhibit a similar sensitivity pattern towards the ΦGP100 phage, while strains BRIP, Cab57 and CMR5c display a moderate to mild sensitivity.

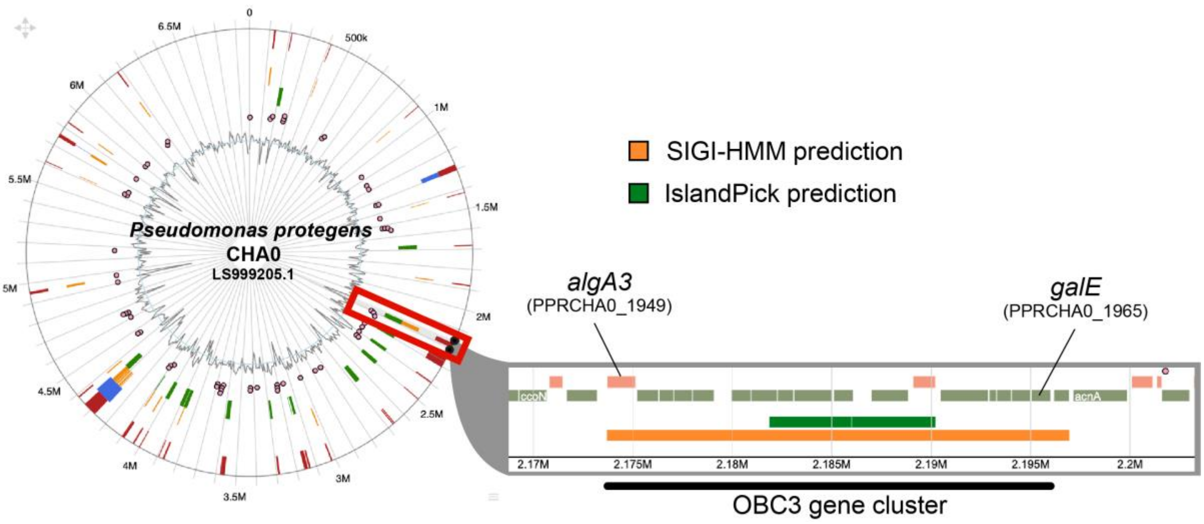


**Fig. S8. The OBC3 gene cluster is located within a potential genomic island.** Predictions were obtained from IslandViewer4 (https://www.pathogenomics.sfu.ca/islandviewer/). The location of the OBC3 gene cluster is highlighted with a red square.

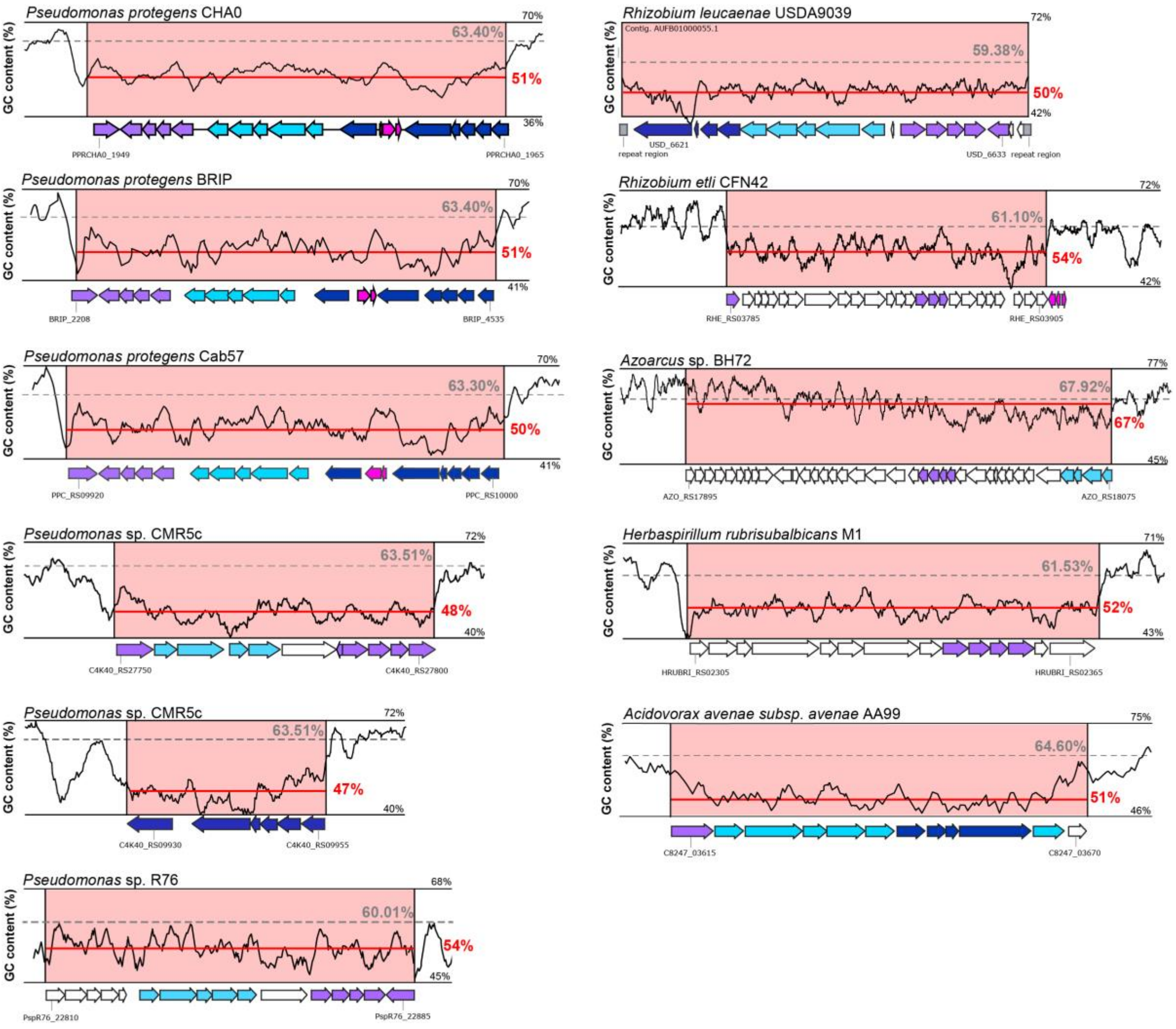


**Fig. S9. Horizontal gene transfer signature detected within the different orthologous genomic regions of OBC3 in bacterial genomes**. The gray dashed lines represent the average GC content of the whole genome, while the red lines correspond to the GC content of the orthologous genomic regions of the OBC3 gene cluster. Dark lines illustrate the GC skew calculated using a 500-bp window frame.

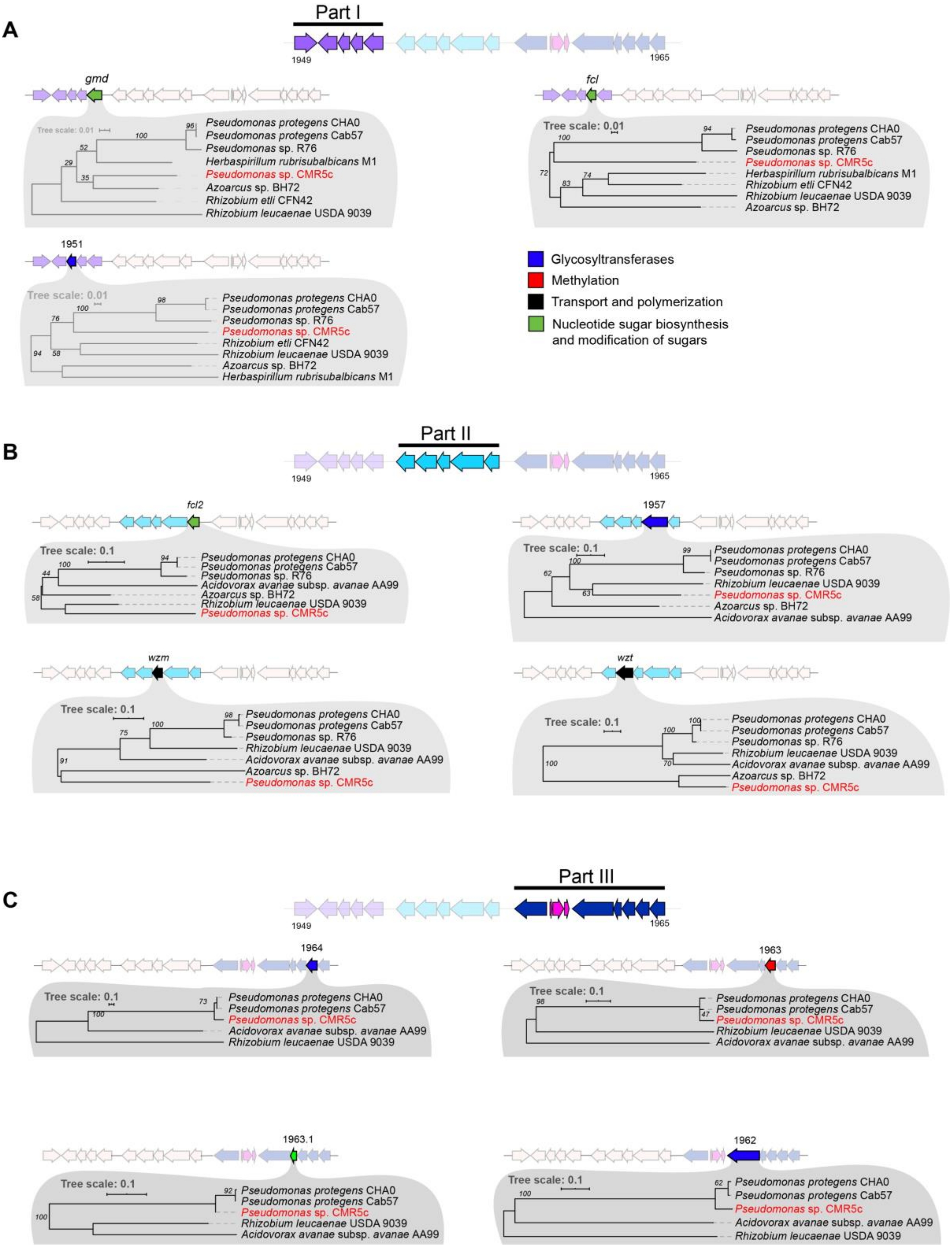


**Fig. S10. Maximum likelihood trees of the amino acid sequences of the corresponding genes conserved within the OBC3 gene cluster detected in bacterial genomes.** Maximum likelihood trees obtained from the aligned amino acid sequences present within the three parts of the OBC3 gene cluster (**A**: part 1; **B**: part 2; **C**: part 3). The robustness of each tree was assessed with 100 bootstraps replicates. *Pseudomonas* strains with names written in red exhibit incongruences in the tree topology compared to the species phylogenetic tree shown in **Fig. 4**.

### Supplementary tables

**Table S1**. Plasmids used in this study.

| Plasmid | Genotype or relevant characteristics[1] | Reference or source |
|---|---|---|
| pEMG | Expression vector; *ori*R6K, *lacZα* with two flanking I-SceI sites; $Km^R$, $Ap^R$ | (2) |
| pRL27 | Tn5-RL27 ($Km^R$-*ori*R6 K) delivery vector for the generation of transposon mutant library. | (3) |
| pME8300 | Carrier plasmid for Tn*7* for $P_{tac/lacIq}$ controlled target gene expression; $Gm^r$, $Ap^r$ | (4) |
| pUX-BF13 | Helper plasmid encoding Tn*7* transposition functions; R6K-replicon; $Ap^r$ | (4) |
| pME497 | Mobilizing plasmid; IncP-1 Tra RepA (Ts); $Ap^R$ | (5) |
| pME11021 | pME8300-$P_{tac/lacIq}$-*fcl*(CFN42); IPTG-inducible expression of *Rhizobium etli* CFN42 *fcl* (RHE_CH00763); $Gm^r$, $Ap^r$ | This study |
| pME11022 | pME8300-$P_{tac/lacIq}$-*fcl*(CHA0); IPTG-inducible expression of *Pseudomonas protegens* CHA0 *fcl*; $Gm^r$, $Ap^r$ | This study |
| pME11107 | pEMG::*Δfcl2*; suicide plasmid for the deletion of the *fcl2*; $Km^R$ | This study |
| pME11108 | pEMG::*Δ1962*; suicide plasmid for the deletion of PPRCHA0_1962; $Km^R$ | This study |
| pME11106 | pEMG::*ΔgalE*; suicide plasmid for the deletion of the *galE* gene; $Km^R$ | This study |
| pME11105 | pEMG::*ΔoatA*; suicide plasmid for the deletion of the *oatA* gene; $Km^R$ | This study |
| pME11123 | pEMG::*algC;* suicide plasmid for the deletion of the *algC* gene; $Km^R$ | This study |

[1] $Ap^R$, ampicillin resistance; $Cm^R$, chloramphenicol resistance; $Gm^R$, gentamycin resistance; $Km^R$, kanamycin resistance; $Tc^R$, tetracycline resistance.

**Table S2.** Tn-seq characteristics.

| Sample | Biological replicate | Total number of reads obtained | Number of HQ reads [a] | Total of mapping HQ reads [b] | Mean read length (bp) | Total insertion hits | Number of insertions every 1,000 bp [c] | Total genes with hits | Mean of hits per genes |
|---|---|---|---|---|---|---|---|---|---|
| Control | 1 | 20,242,678 | 11,731,031 | 9,769,299 (83%) | 113.3 | 526,158 | 77 | 5,788 (94%) | 82 |
| Control | 2 | 22,399,901 | 13,419,248 | 11,237,426 (84%) | 112.6 | 425,745 | 63 | 5,699 (93%) | 67 |
| Control | 3 | 22,967,708 | 13,804,616 | 11,411,687 (83%) | 113.2 | 546,931 | 77 | 5,854 (95%) | 84 |
| MOI of 1 [d] | 1 | 26,759,802 | 11,170,004 | 9,282,169 (83%) | 112.9 | 55,159 | - | 4,888 (79%) | 10 |
| MOI of 1 | 2 | 22,880,585 | 9,139,588 | 7,694,614 (84%) | 113.6 | 75,459 | - | 5,681 (92%) | 12 |
| MOI of 1 | 3 | 23,263,048 | 13,171,418 | 11,238,630 (85%) | 113.9 | 52,282 | - | 4,649 (76%) | 10 |
| MOI of 10 | 1 | 20,814,448 | 11,854,767 | 9,855,684 (83%) | 112.9 | 46,562 | - | 4,605 (75%) | 9 |
| MOI of 10 | 2 | 20,297,964 | 10,598,935 | 8,989,953 (85%) | 112.5 | 49,067 | - | 4,815 (78%) | 9 |
| MOI of 10 | 3 | 16,808,770 | 9,302,312 | 7,851,007 (84%) | 113.5 | 42,104 | - | 4,425 (72%) | 9 |
| MOI of 100 | 1 | 20,665,244 | 9,817,868 | 8,286,692 (84%) | 112.5 | 44,272 | - | 4,581 (74%) | 9 |
| MOI of 100 | 2 | 18,191,640 | 10,076,908 | 8,619,761 (86%) | 114.1 | 44,946 | - | 4,570 (74%) | 9 |
| MOI of 100 | 3 | 21,690,076 | 12,089,051 | 10,315,697 (85%) | 113.9 | 46,204 | - | 4,760 (77%) | 9 |

[a] HQ reads, high-quality reads (i.e., trimmed reads and reads that matched with the Tn*5* transposon sequence).

[b] Number of high-quality reads that mapped to the genome of CHA0 (LS999205.1).

[c] Total number of genes in the genome of *P. protegens* CHA0: 6,158.

[d] MOI, multiplicity of infection.

**Table S3.** Phage genomes used in this study.

| Accession Number | Taxid | Genome length (bp) | Name of the phage | Type | Phage family | Bacterial host group | Bacterial host name |
|---|---|---|---|---|---|---|---|
| NC_003907 | 2907796 | 46,012 | *Vibrio* phage VpV262 | dsDNA | *Zobellviridae* | Gammaproteobacteria | *Vibrio parahaemolyticus* |
| NC_048671 | 2847823 | 40,907 | *Lentibacter* phage vB_LenP_ICBM2 | dsDNA | *Zobellviridae* | Alphaproteobacteria | *Lentibacter sp.* |
| NC_018280 | 1197951 | 38,889 | *Celeribacter* phage P12053L | dsDNA | *Zobellviridae* | Alphaproteobacteria | *Celeribacter marinus* |
| NC_002519 | 2905867 | 39,898 | *Roseobacter* phage SIO1 | dsDNA | *Zobellviridae* | Alphaproteobacteria | *Roseobacter* |
| NC_048672 | 2847822 | 40,163 | *Lentibacter* phage vB_LenP_ICBM1 | dsDNA | *Zobellviridae* | Alphaproteobacteria | *Lentibacter sp.* |
| NC_024791 | 1529058 | 50,440 | *Vibrio* phage ICP2_2013_A_Haiti | dsDNA | *Zobellviridae* | Gammaproteobacteria | *Vibrio cholerae* |
| NC_015158 | 979533 | 49,675 | *Vibrio* phage ICP2 | dsDNA | *Zobellviridae* | Gammaproteobacteria | *Vibrio cholerae* |
| KM224878 | 1529057 | 50,250 | *Vibrio* phage ICP2_2011_A | dsDNA | *Zobellviridae* | Gammaproteobacteria | *Vibrio cholerae* |
| HQ641346 | 979534 | 48,626 | *Vibrio* phage ICP2_2006_A | dsDNA | *Zobellviridae* | Gammaproteobacteria | *Vibrio cholerae* |
| NC_021300 | 1316739 | 43,882 | *Pseudoalteromonas* phage RIO-1 | dsDNA | *Zobellviridae* | Gammaproteobacteria | *Pseudoalteromonas marina* |
| NC_048630 | 1357706 | 45,035 | *Pseudoalteromonas* phage HP1 | dsDNA | *Zobellviridae* | Gammaproteobacteria | *Pseudoalteromonas sp.* |
| NC_019540 | 1161935 | 49,390 | *Salinivibrio* phage CW02 | dsDNA | *Zobellviridae* | Gammaproteobacteria | *Salinivibrio costicola* |
| NC_027988 | 1622234 | 47,636 | *Citrobacter* phage CVT22 | dsDNA | *Zobellviridae* | Gammaproteobacteria | *Citrobacter sp.* TM1552 |
| LT986460 | 2055238 | 50,547 | *Pseudomonas* phage GP100 | dsDNA | *Zobellviridae* | Gammaproteobacteria | *Pseudomonas protegens* |
| NC_007808 | 347327 | 49,639 | *Pseudomonas* phage PA11 | dsDNA | *Zobellviridae* | Gammaproteobacteria | *Pseudomonas aeruginosa* |
| NC_031274 | 1784982 | 50,509 | *Pseudomonas* phage O4 | dsDNA | *Zobellviridae* | Gammaproteobacteria | *Pseudomonas aeruginosa* PAO1 |
| NC_018859 | 1604355 | 77,315 | *Escherichia* phage ECBP2 | dsDNA | - | Gammaproteobacteria | *Escherichia coli* |
| NC_023593 | 1436889 | 76,184 | *Escherichia* phage KBNP1711 | dsDNA | - | Gammaproteobacteria | *Escherichia coli* |
| NC_027395 | 1519788 | 77,327 | *Escherichia* phage vB_EcoP_SU10 | dsDNA | - | Gammaproteobacteria | *Escherichia coli* |
| NC_010324 | 2679905 | 77,554 | *Escherichia* phage phiEco32 | dsDNA | - | Gammaproteobacteria | *Escherichia coli* |
| NC_018835 | 1237159 | 77,448 | *Escherichia* phage NJ01 | dsDNA | - | Gammaproteobacteria | *Escherichia coli* |
| NC_028903 | 1598146 | 77,266 | *Escherichia* phage 172-1 | dsDNA | - | Gammaproteobacteria | *Escherichia coli* |
| NC_015938 | 1054968 | 89,916 | *Salmonella* phage 7-11 | dsDNA | *Podoviridae* | Gammaproteobacteria | *Salmonella enterica subsp. enterica serovar Newport* |
| NC_019402 | 1141137 | 76,631 | *Cronobacter* phage vB_CsaP_GAP52 | dsDNA | *Podoviridae* | Gammaproteobacteria | *Cronobacter sakazakii* |
| NC_048861 | 2712958 | 90,710 | *Proteus* phage Privateer | dsDNA | - | Gammaproteobacteria | *Proteus mirabilis* |
| NC_048664 | 2699738 | 92,122 | *Cronobacter* phage vB_CsaP_009 | dsDNA | - | Gammaproteobacteria | *Cronobacter sakazakii* |

| | | | | | | | |
|---|---|---|---|---|---|---|---|
| NC_048776 | 2591033 | 97,988 | *Aeromonas* phage LAh_9 | dsDNA | - | Gammaproteobacteria | *Aeromonas hydrophila* |
| NC_048773 | 2588517 | 102,915 | *Aeromonas* phage 4_4572 | dsDNA | - | Gammaproteobacteria | *Aeromonas hydrophila subsp. hydrophila* |
| NC_048775 | 2591032 | 97,408 | *Aeromonas* phage LAh_8 | dsDNA | - | Gammaproteobacteria | *Aeromonas hydrophila* |
| NC_048774 | 2591030 | 101,390 | *Aeromonas* phage LAh_6 | dsDNA | - | Gammaproteobacteria | *Aeromonas hydrophila* |
| KJ936628 | 1524880 | 50,431 | *Vibrio* phage VPp1 | dsDNA | *Podoviridae* | Gammaproteobacteria | *Vibrio parahaemolyticus* |
| NC_042103 | 2079288 | 45,936 | *Pseudomonas* phage Bjorn | dsDNA | - | Gammaproteobacteria | *Pseudomonas* sp. |
| NC_017971 | 1114179 | 46,271 | *Pseudomonas* phage tf | dsDNA | - | Gammaproteobacteria | *Pseudomonas putida* |
| NC_018850 | 1235661 | 45,517 | *Pseudomonas* phage UFV-P2 | dsDNA | - | Gammaproteobacteria | *Pseudomonas fluorescens* |
| NC_042107 | 2079543 | 45,058 | *Pseudomonas* phage NV1 | dsDNA | - | Gammaproteobacteria | *Pseudomonas tolaasii* NCPPB 2192 |
| NC_028933 | 1589273 | 45,626 | *Pseudomonas* phage PhiCHU | dsDNA | - | Gammaproteobacteria | *Pseudomonas aeruginosa* |
| NC_004466 | 2905964 | 45,503 | *Pseudomonas* phage PaP3 | dsDNA | - | Gammaproteobacteria | *Pseudomonas* |
| JN254801 | 1158721 | 44,789 | *Pseudomonas* phage MR299-2 | dsDNA | - | Gammaproteobacteria | *Pseudomonas aeruginosa* |
| NC_019813 | 1234701 | 44,030 | *Pseudomonas* phage vB_PaeP_p2-10_Or1 | dsDNA | - | Gammaproteobacteria | *Pseudomonas aeruginosa* |
| HE983844 | 1229675 | 45,469 | *Pseudomonas* phage vB_PaeP_C1-14_Or | dsDNA | - | Gammaproteobacteria | *Pseudomonas aeruginosa* |
| NC_026599 | 1548906 | 45,808 | *Pseudomonas* phage vB_PaeP_C2-10_Ab22 | dsDNA | - | Gammaproteobacteria | *Pseudomonas aeruginosa* |
| NC_023583 | 1406974 | 45,696 | *Pseudomonas* phage TL | dsDNA | - | Gammaproteobacteria | *Pseudomonas aeruginosa* |
| NC_010325 | 484895 | 45,625 | *Bruynoghevirus* LUZ24 | dsDNA | - | Gammaproteobacteria | *Pseudomonas aeruginosa* |
| NC_042343 | 1273709 | 43,895 | *Pseudomonas* phage PaP4 | dsDNA | - | Gammaproteobacteria | *Pseudomonas aeruginosa* |
| NC_022971 | 1429758 | 45,344 | *Pseudomonas* phage phiIBB-PAA2 | dsDNA | - | Gammaproteobacteria | *Pseudomonas aeruginosa* |
| NC_028919 | 1640969 | 45,673 | *Pseudomonas* phage DL54 | dsDNA | - | Gammaproteobacteria | *Pseudomonas aeruginosa* PAO1 |

**Table S4.** Update of the phage ΦGP100 annotation.

| Locus_tag | Start | End | Size (bp) | Strand | Predicted function [Associated Source Organism] | Accession | Identity % |
|---|---|---|---|---|---|---|---|
| GP100_00001 | 424 | 573 | 149 | + | Hypothetical protein | | |
| GP100_00002 | 573 | 644 | 71 | + | Hammerhead_II | | |
| GP100_00003 | 1570 | 1833 | 263 | + | Hypothetical protein | | |
| GP100_00004 | 2034 | 2106 | 72 | + | tRNA-Stop(tta) | | |
| GP100_00005 | 2129 | 2689 | 560 | + | Coil containing protein [*Vibrio* phage 1.204.O._10N.222.46.F12] | YP_009817597.1 | 43/110 (39%) |
| GP100_00006 | 2820 | 2966 | 146 | + | Hypothetical protein | | |
| GP100_00007 | 2966 | 3163 | 197 | + | Hypothetical protein | | |
| GP100_00008 | 3391 | 3729 | 338 | + | Hypothetical protein | | |
| GP100_00009 | 3779 | 3940 | 161 | + | Hypothetical protein | | |
| GP100_00010 | 4265 | 4480 | 215 | + | Hypothetical protein | | |
| GP100_00011 | 4489 | 4779 | 290 | + | Hypothetical protein | | |
| GP100_00012 | 4926 | 5072 | 146 | + | Hypothetical protein | | |
| GP100_00013 | 5072 | 5392 | 320 | + | Hypothetical protein | | |
| GP100_00014 | 5593 | 5928 | 335 | + | Hypothetical protein | | |
| GP100_00015 | 5978 | 6175 | 197 | + | Hypothetical protein | | |
| GP100_00016 | 6148 | 6288 | 140 | - | Hypothetical protein | | |
| GP100_00017 | 6315 | 6449 | 134 | + | Hypothetical protein | | |
| GP100_00018 | 6449 | 6634 | 185 | + | Hypothetical protein | | |
| GP100_00019 | 6727 | 6930 | 203 | + | Hypothetical protein | | |
| GP100_00020 | 6923 | 7876 | 953 | + | Hypothetical protein | | |
| GP100_00021 | 7869 | 8504 | 635 | + | Putative amidotransferase [*Pseudomonas* phage O4] | YP_009304537.1 | 116/213 (54%) |
| GP100_00022 | 8501 | 8719 | 218 | + | Hypothetical protein | | |
| GP100_00023 | 8774 | 10147 | 1373 | + | Putative amidotransferase [*Pseudomonas* phage O4] | YP_009304535.1 | 158/415 (38%) |
| GP100_00024 | 10180 | 10647 | 467 | + | Putative HNH endonuclease [*Escherichia* phage LL11] | YP_009812380.1 | 60/148 (41%) |
| GP100_00025 | 10637 | 11506 | 869 | + | Putative COOH-NH2 ligase [*Pseudomonas* phage O4] | YP_009304534.1 | 168/275 (61%) |

| | | | | | | | |
|---|---|---|---|---|---|---|---|
| GP100_00026 | 11508 | 12065 | 557 | + | Hypothetical protein | | |
| GP100_00027 | 12062 | 12946 | 884 | + | Putative amidoligase [*Pseudomonas* phage O4] | YP_009304532.1 | 169/275 (61%) |
| GP100_00028 | 12946 | 13155 | 209 | + | Hypothetical protein | | |
| GP100_00029 | 13152 | 14285 | 1133 | + | Putative ATP-grasp protein [*Pseudomonas* phage O4] | YP_009304531.1 | 141/301 (47%) |
| GP100_00030 | 14353 | 14565 | 212 | + | Hypothetical protein | | |
| GP100_00031 | 14574 | 14750 | 176 | + | Hypothetical protein | | |
| GP100_00032 | 14762 | 16483 | 1721 | + | Putative helicase/primase [*Pseudomonas* phage O4] | YP_009304525.1 | 398/570 (70%) |
| GP100_00033 | 16483 | 16650 | 167 | + | Recombination protein RecR | | |
| GP100_00034 | 16637 | 17125 | 488 | + | RNA polymerase sigma factor [*Pseudomonas* phage O4] | YP_009304524.1 | 73/159 (46%) |
| GP100_00035 | 17115 | 18992 | 1877 | + | Putative DNA polymerase [*Pseudomonas* phage O4] | YP_009304521.1 | 389/625 (62%) |
| GP100_00036 | 18976 | 19077 | 101 | + | Hypothetical protein | | |
| GP100_00037 | 19077 | 19430 | 353 | + | Hypothetical protein | | |
| GP100_00038 | 19434 | 20141 | 707 | + | Putative ssDNA binding protein [*Pseudoalteromonas* phage RIO-1] | YP_008051092.1 | 65/193 (34%) |
| GP100_00039 | 20135 | 20320 | 185 | + | Hypothetical protein | | |
| GP100_00040 | 20392 | 20766 | 374 | + | Hypothetical protein | | |
| GP100_00041 | 20757 | 20942 | 185 | + | Hypothetical protein | | |
| GP100_00042 | 20926 | 21804 | 878 | + | Putative 5'-3' exonuclease [*Pseudomonas* phage O4] | YP_009304516.1 | 166/286 (58%) |
| GP100_00043 | 21797 | 22336 | 539 | + | Metallophosphoesterase [*Ralstonia* phage RSP15] | YP_009277057.1 | 82/181 (45%) |
| GP100_00044 | 22336 | 22869 | 533 | + | Hypothetical protein | | |
| GP100_00045 | 23040 | 23192 | 152 | + | Hypothetical protein | | |
| GP100_00046 | 23176 | 23619 | 443 | + | Putative terminase small subunit [*Pseudomonas* phage O4] | YP_009304514.1 | 89/134 (66%) |
| GP100_00047 | 23606 | 23860 | 254 | + | Hypothetical protein | | |
| GP100_00048 | 23857 | 24192 | 335 | + | Putative AAT_I superfamily protein [*Pectobacterium* phage PPWS2] | YP_009816196.1 | 34/88 (39%) |
| GP100_00049 | 24179 | 24952 | 773 | + | Putative 3'-5' exonuclease [*Pseudomonas* phage O4] | YP_009304511.1 | 193/252 (77%) |
| GP100_00050 | 25166 | 25534 | 368 | + | Putative holin [*Pseudomonas* phage UFV-P2] | YP_007518479.1 | 64/87 (74%) |
| GP100_00051 | 25620 | 26225 | 605 | + | PhoH-like protein [*Pseudomonas* phage PA11] | YP_001294617.1 | 155/195 (79%) |
| GP100_00052 | 26225 | 26527 | 302 | + | Hypothetical protein | | |
| GP100_00053 | 26521 | 26670 | 149 | + | Hypothetical protein | | |
| GP100_00054 | 26667 | 26861 | 194 | + | Hypothetical protein | | |

| | | | | | | | |
|---|---|---|---|---|---|---|---|
| GP100_00055 | 26900 | 27163 | 263 | + | Neck appendage protein precursor [*Bacillus* phage GA1] | NP_073695 | 17/65 (26%) |
| GP100_00056 | 27199 | 27483 | 284 | - | Hypothetical protein | | |
| GP100_00057 | 27576 | 28049 | 473 | - | Hypothetical protein | | |
| GP100_00058 | 28027 | 28266 | 239 | - | Hypothetical protein | | |
| GP100_00059 | 28269 | 28703 | 434 | - | Putative endolysin [*Pseudomonas* phage O4] | YP_009304499.1 | 118/143 (83%) |
| GP100_00060 | 28703 | 28807 | 104 | - | Hypothetical protein | | |
| GP100_00061 | 28809 | 30641 | 1832 | - | Putative IVP-C-like protein [*Klebsiella* phage vB_KpnP_Klyazma] | UVD32011.1 | 177/613 (29%) |
| GP100_00062 | 30643 | 35430 | 4787 | - | Putative IVP-D-like protein [*Klebsiella* phage vB_KpnP_Klyazma] | UVD32010.1 | 520/1652 (31%) |
| GP100_00063 | 35443 | 35964 | 521 | - | Hypothetical protein | | |
| GP100_00064 | 35969 | 36415 | 446 | - | Acyl-CoA N-acyltransferase (GNAT) domain protein [*Lentibacter* virus vB_LenP_ICBM1] | YP_009834282.1 | 26/84 (31%) |
| GP100_00065 | 36408 | 38705 | 2297 | - | Putative head closure protein [*Pseudomonas* phage O4] | YP_009304494.1 | 407/772 (53%) |
| GP100_00066 | 38689 | 38871 | 182 | - | Hypothetical protein | | |
| GP100_00067 | 38871 | 39677 | 806 | - | Putative adaptor protein [*Pseudomonas* phage O4] | YP_009304492.1 | 152/259 (59%) |
| GP100_00068 | 39677 | 40303 | 626 | - | Neck protein [*Raoultella* phage RP180] | YP_009821849.1 | 58/151 (38%) |
| GP100_00069 | 40408 | 41052 | 644 | - | Putative tail shaft protein Tsh [*Listeria* phage vB_LmoS_188] | YP_009204855.1 | 40/94 (43%) |
| GP100_00070 | 41068 | 42021 | 953 | - | Putative capsid protein [*Pseudomonas* phage O4] | YP_009304488.1 | 234/311 (75%) |
| GP100_00071 | 42034 | 42792 | 758 | - | Putative scaffold protein [*Pseudomonas* phage O4] | YP_009304487.1 | 119/252 (47%) |
| GP100_00072 | 42902 | 42976 | 74 | - | tRNA-Asn(gtt) | | |
| GP100_00073 | 43104 | 43343 | 239 | - | Hypothetical protein | | |
| GP100_00074 | 43346 | 45100 | 1754 | - | Putative portal protein [*Pseudomonas* phage O4] | YP_009304485.1 | 422/584 (72%) |
| GP100_00075 | 45110 | 46714 | 1604 | - | Putative terminase large subunit [*Pseudomonas* phage O4] | YP_009304484.1 | 397/533 (74%) |
| GP100_00076 | 46714 | 47214 | 500 | - | Hypothetical protein | | |
| GP100_00077 | 47250 | 48761 | 1511 | - | Putative pectate lyase (potential depolymerase) | | |
| GP100_00078 | 48761 | 49684 | 923 | - | Hypothetical protein | | |
| GP100_00079 | 49681 | 50328 | 647 | - | Hypothetical protein | | |

**Table S5.** Bacterial strains used in this study.

| Strain [a] | Genome accession no. | Origin | Reference or source |
|---|---|---|---|
| ***Pseudomonas protegens* subgroup** | | | |
| *P. protegens* BRIP | NZ_LHUW00000000.1 | Cyclops | (6, 7) |
| *P. protegens* Cab57 | NZ_AP014522.1 | Shepperd's purse rhizosphere | (8) |
| *P. protegens* CHA0$^T$ | LS999205.1 | Tobacco rhizosphere | (9, 10) |
| *P. protegens* K94.41 | NZ_LHUU00000000.1 | Cucumber rhizosphere | (7, 11) |
| *P. protegens* Pf-5 | NC_004129.6 | Cotton rhizosphere | (12, 13) |
| *P. protegens* PF | NZ_LHUX00000000.1 | Wheat leaves | (7, 14) |
| *P. protegens* Pf1 | ERS3935804 | Tobacco | (14) |
| *P. protegens* PGNR1 | NZ_LHUV00000000.1 | Tobacco rhizosphere | (7, 14) |
| *P. saponiphila* DMS9751$^T$ | FNTJ00000000.1 | unknown | (15) |
| *Pseudomonas* sp. AU11706 | NZ_LCZB00000000.1 | Cystic fibrosis sputum | (16) |
| *Pseudomonas* sp. AU20219 | NZ_LDET00000000.1 | Cystic fibrosis sputum | (16) |
| *Pseudomonas* sp. AU13852 | NZ_LCZC00000000.1 | Cystic fibrosis sputum | (16) |
| *Pseudomonas* sp. CMR5c | NZ_LHUY00000000.1 | Red cocoyam rhizosphere | (7, 17) |
| *Pseudomonas* sp. CMR12a | CP027706.1 | Red cocoyam rhizosphere | (7, 17) |
| *Pseudomonas* sp. Pf4 | LUUD00000000.1 | Lamb's lettuce | (18) |
| *Pseudomonas* sp. Pf11 | LUUE00000000.1 | Lamb's lettuce | (18) |
| *Pseudomonas* sp. LD120 | ERR3588830 | Blade of marine alga | (19, 20) |
| *Pseudomonas* sp. Os17 | NZ_AP014627.1 | Rice rhizosphere | (8) |
| *Pseudomonas* sp. St29 | NZ_AP014628.1 | Potato rhizosphere | (8) |
| ***Pseudomonas chlororaphis* subgroup** | | | |
| *P. chlororaphis* O6 | NZ_CM001490.1 | Soil | (21) |
| *P. chlororaphis* R47 | Genome not published | Potato rhizosphere | (22) |
| *P. chlororaphis* subsp. *aureofaciens* 30-84 | NZ_CM001559.1 | Wheat rhizosphere | (21, 23) |
| *P. chlororaphis* subsp. *aurantiaca* JD37 | NZ_CP009290.1 | Potato rhizosphere | (24, 25) |
| *P. chlororaphis* subsp. *aureofaciens* CD | NZ_LHVB00000000.1 | Cyclops (water) | (6, 7) |
| *P. chlororaphis* subsp. *aureofaciens* LMG1245$^T$ | NZ_LHVA00000000.1 | River clay | (7, 26) |
| *P. chlororaphis* subsp. *chlororaphis* LMG5004$^T$ | NZ_LHVC00000000.1 | Contaminated plate | (7, 27) |
| *P. chlororaphis* subsp. *piscium* DSM21509$^T$ | NZ_CP027707.1 | European perch intestine | (28) |
| *P. chlororaphis* subsp. *piscium* PCL1391 | NZ_CP027736.1 | Tomato root | (7, 29) |
| **Other species** | | | |
| *Acidovorax avenae* subsp. *avenae* AA99 | CP028289.1 | Maize | (30) |
| *Azoarcus* sp. BH72 | NC_008702.1 | Kallar grass | (31) |
| *Herbaspirillum rubrisubalbicans* M1 | CP013737.1 | Sugar cane | (32) |
| *Rhizobium etli* CFN42$^T$ | CP000133.1 | Common bean | (33) |
| *Rhizobium leucaenae* USDA 9039$^T$ | AUFB01000000.1 | *Leucaena esculenta* | (34) |

[a] $^T$, type strain

**Table S6.** Occurrence of inverted repeats sequences inside the OBC3 gene cluster of *Pseudomonas protegens* CHA0.

| Inverted repeat | Stem | Loop | Sequence | Occurrence inside the genome of CHA0 |
|---|---|---|---|---|
| IR1 | 17 | 9,928 | 5'-CGCTTTTCAAACTCTAC-3'<br>3'-GCGAAAAGTTTGAGATG-5' | Unique |
| IR2 | 13 | 15,271 | 5'-AGCTGAAGGCAGT-3'<br>3'-TCGACTTCCGTCA-5' | Unique |
| IR3 | 13 | 7,086 | 5'-GCAGGCGACGCAA-3'<br>3'-CGTCCGCTGCGTT-5' | Unique |
| IR4 | 13 | 4,123 | 5'-TGCGAAAGACTAC-3'<br>3'-ACGCTTTCTGATG-5' | Unique |
| IR5 | 13 | 7,221 | 5'-TTGCATCGCTTGG-3'<br>3'-AACGTAGCGAACC-5' | Unique |
| IR6 | 12 | 3,406 | 5'-ATTAGCGGCGCA-3'<br>3'-TAATCGCCGCGT-5' | Unique |
| IR7 | 12 | 13,520 | 5'-CTAACTATTACT-3'<br>3'-GATTGATAATGA-5' | Unique |
| IR8 | 12 | 2,975 | 5'-GCTTTGCTACTG-3'<br>3'-CGAAACGATGAC-5' | Unique |
| IR9 | 12 | 8,540 | 5'-TTTGGTCGCTTC-3'<br>3'-AAACCAGCGAAG-5' | Unique |
| IR10 | 12 | 15,176 | 5'-TTTTGGCAGCCC-3'<br>3'-AAAACCGTCGGG-5' | Three times |
| IR11 | 12 | 8,568 | 5'-ACGCCCTGAAAA-3'<br>3'-TGCGGGACTTTT-5' | Four times |
| IR12 | 12 | 4 | 5'-ATCCAGCGGCGG-3'<br>3'-TAGGTCGCCGCC-5' | Six times |
| IR13 | 12 | 5,410 | 5'-TGAAGCTGGCAG-3'<br>3'-ACTTCGACCGTC-5' | Six times |
| IR14 | 12 | 11,731 | 5'-TTCATTGACCAG-3'<br>3'-AAGTAACTGGTC-5' | Seven times |

**Table S7**. *Pseudomonas protegens* CHA0 derivatives and *Escherichia coli* strain used in this study.

| Strain name | Strain code | Genotype or relevant characteristics[a] | Reference or source |
|---|---|---|---|
| ***Pseudomonas protegens*** **CHA0 and derivatives** | | | |
| CHA0$^T$ | CHA0$^T$ | *P. protegens* type strain; wild type; genome accession no. LS999205.1 | (9, 10) |
| Δ*fcl* | CHA5205 | CHA0 with deletion of PPRCHA0_1952 | (35) |
| Δ*wbpL* | CHA5161 | CHA0 with deletion of PPRCHA0_4350 | (35) |
| Δ*wzx* | CHA5206 | CHA0 with deletion of PPRCHA0_4354 | (35) |
| Δ*fcl2* | CHA5328 | CHA0 with deletion of PPRCHA0_1958 | This study |
| Δ*galE* | CHA5327 | CHA0 with deletion of PPRCHA0_1965 | This study |
| Δ*1962* | CHA5330 | CHA0 with deletion of PPRCHA0_1962 | This study |
| Δobc3 | CHA5182 | CHA0 with deletion of PPRCHA0_1950 through PPRCHA0_1958 | (35) |
| Δ*oatA* | CHA5329 | CHA0 with deletion of PPRCHA0_1960 | This study |
| Δ*algC* | CHA5303 | CHA0 with deletion of PPRCHA0_5962 | This study |
| CHA5211 | CHA5211 | CHA5205::attTn*7*-$P_{tac/lacIq}$-*fcl*(CFN42); Gm$^r$ | This study |
| CHA5212 | CHA5212 | CHA5205::attTn*7*-$P_{tac/lacIq}$-*fcl*(CHA0); Gm$^r$ | This study |
| ***Escherichia coli*** | | | |
| *E. coli* S17-1/λpir | | Laboratory strain | (36) |

[a] Gm$^R$, gentamicin resistance.

**Table S8**. Primers used in this study.

| Name | Sequence 5' → 3', restriction enzyme(s)[a] | Purpose |
|---|---|---|
| del-fcl2-1 | CGGAATTCTACCATTCGGGCCACATG, EcoRI | Deletion of *fcl2* (PPRCHA0_1958) |
| del-fcl2-2 | GCTCTAGAATCTGATGCAAGCACCGC, XbaI | Deletion of *fcl2* (PPRCHA0_1958) |
| del-fcl2-3 | GCTCTAGACATTCCTTTGTGCCCTGT, XbaI | Deletion of *fcl2* (PPRCHA0_1958) |
| del-fcl2-4 | ACGCGTCGACATGTAACGCAGCACCTCT, SalI | Deletion of *fcl2* (PPRCHA0_1958) |
| fcl2-check-F | GAAGTGTGCGGGTAAGTG | Control of deletion of *fcl2* (PPRCHA0_1958) |
| fcl2-check-R | AATGTGCCAATGCCCTTG | Control of deletion of *fcl2* (PPRCHA0_1958) |
| del-algC-1 | ggGGTACCATGCACGCTTTACAAGCC, KpnI | Deletion of *algC* (PPRCHA0_5962) |
| del-algC-2 | cgGGATCCCATCACTGAGGGTGCTCC, BamHI | Deletion of *algC* (PPRCHA0_5962) |
| del-algC-3 | gGGATCCTGATTGTTTGAAGCACCG, BamHI | Deletion of *algC* (PPRCHA0_5962) |
| del-algC-4 | acgcGTCGACGATGTTGTAGGACTCGCC, SalI | Deletion of *algC* (PPRCHA0_5962) |
| algC-check-F | GCCAGCTGATCTCTTTTATC | Control of deletion of *algC* (PPRCHA0_5962) |
| algC-check-R | TTTTCGGCTTTCAATGCTTC | Control of deletion of *algC* (PPRCHA0_5962) |
| del-oatA-1 | CGGAATTCCGATGTTACCACGTCGTA, EcoRI | Deletion of *oatA* (PPRCHA0_1959) |
| del-oatA-2 | CGAGCTCTAACGCCTCAATACGTTATCA, SacI | Deletion of *oatA* (PPRCHA0_1959) |
| del-oatA-3 | CGAGCTCCATGTCTTGGAAAAATCCTTG, SacI | Deletion of *oatA* (PPRCHA0_1959) |
| del-oatA-4 | GGGGTACCTCCTTGGCAAAGTATGCG, KpnI | Deletion of *oatA* (PPRCHA0_1959) |
| oatA-check-F | TTCCTGTAGCCCTTCTGC | Control of deletion of *oatA* (PPRCHA0_1959) |
| oatA-check-R | GCGCGCAGCTTGATAAAG | Control of deletion of *oatA* (PPRCHA0_1959) |
| del-1962-1 | CGGAATTCAATTTGATTGCCAGCATGAGT, EcoRI | Deletion of PPRCHA0_1962 |
| del-1962-2 | GGGGTACCCTTTGATGGCGAGCGTTG, KpnI | Deletion of PPRCHA0_1962 |
| del-1962-3 | GGGGTACCCATCCTGGAATGTACCTTGTG, KpnI | Deletion of PPRCHA0_1962 |
| del-1962-4 | ACGCGTCGACATGATTGCACTGGCCATT, SalI | Deletion of PPRCHA0_1962 |
| 1962-check-F | CAGTACAGCAGCTCCGAT | Control of deletion of PPRCHA0_1962 |
| 1962-check-R | CGCACTAGGTGCTCAGTA | Control of deletion of PPRCHA0_1962 |
| del-galE-1 | CGGAATTCATCTGCAGGGATGTCGAT, EcoRI | Deletion of *galE* (PPRCHA0_1965) |
| del-galE-2 | GGGGTACCTAACGCAAATGACGCAATCTT, KpnI | Deletion of *galE* (PPRCHA0_1965) |
| del-galE-3 | GGGGTACCCACACCGCCTCTCCTTCG, KpnI | Deletion of *galE* (PPRCHA0_1965) |
| del-galE-4 | ACGCGTCGACTTGACCCTGACGGCAATT, SalI | Deletion of *galE* (PPRCHA0_1965) |
| galE-check-F | CTCGACGACTCATCAGGC | Control of deletion of *galE* (PPRCHA0_1965) |
| galE-check-R | TATGCCCGTGGATTGTTG | Control of deletion of *galE* (PPRCHA0_1965) |
| 8300-F | ACATCATAACGGTTCTGGCAAA | Amplification of pME8300 to check insertions |
| 8300-R | AAAAAGGGGACCTCTAGGGTC | Amplification of pME8300 to check insertions |
| pEMG check-F | GTAAAACGACGGCCAGT | Sequencing verification of the pEMG-based plasmids |
| pEMG check-R | AACAGCTATGACCATG | Sequencing verification of the pEMG-based plasmids |

[a] Restriction sites are underlined.
